# Genomic Modules and Intramodular Network Convergency of Susceptibility and Resilience in Multimodeled Stress in Male Mice

**DOI:** 10.1101/2021.06.30.450390

**Authors:** Jordan Marrocco, Salvatore G. Caradonna, Tie-Yuan Zhang, Nicholas O’Toole, Mo-Jun Shen, Huzefa Khalil, Nathan R. Einhorn, Xianglan Wen, Carine Parent, Francis S. Lee, Huda Akil, Bruce S. McEwen, Michael J. Meaney

**Affiliations:** Laboratory of Neuroendocrinology, The Rockefeller University, New York, NY, US; Douglas Mental Health University Institute, McGill University, Montreal, Canada; Singapore Institute for Clinical Sciences, Singapore; Michigan Neuroscience Institute, University of Michigan, Ann Arbor, MI, US; Department of Psychiatry, Sackler Institute for Developmental Psychobiology, Weill Cornell Medical College, New York, NY, US; Yong Loo Lin School of Medicine, Singapore; Sackler Program for Epigenetics & Psychobiology at McGill University

## Abstract

The multifactorial etiology of stress-related disorders is a challenge in developing synchronized medical standards for treatment and diagnosis. It is largely unknown whether there exists molecular convergence in preclinical models of stress generated using disparate construct validity. Using RNA-sequencing (RNA-seq), we investigated the genomic signatures in the ventral hippocampus, which mostly regulates affective behavior, in mouse models that recapitulate the hallmarks of anxiety and depression. Chronic oral corticosterone (CORT), a model that causes a blunted endocrine response to stress, induced anxiety- and depression-like behavior in wildtype mice and mice heterozygous for the gene coding for brain-derived neurotrophic factor (BDNF) Val66Met, a variant associated with genetic susceptibility to stress. In a separate set of mice, chronic social defeat stress led to a susceptible or a resilient population, whose proportion was dependent on housing conditions, standard housing or enriched environment. A rank-rank-hypergeometric (RRHO) analysis of the RNA-seq data set across models demonstrated that in mice treated with CORT and susceptible mice raised in standard housing differentially expressed genes (DEGs) converged toward gene networks involved in similar biological functions. Weighted gene co-expression analysis generated 54 unique modules of interconnected gene hubs, two of which included a combination of all experimental groups and were significantly enriched in DEGs, whose function was consistent with that predicted in the RRHO GO analysis. This multimodel approach showed transcriptional synchrony between models of stress with hormonal, environmental or genetic construct validity shedding light on common genomic drivers that embody the multifaceted nature of stress-related disorders.

## INTRODUCTION

Stressors induce a variety of molecular and physiological mechanisms, influenced by genes and environment, leading to alterations in coping mechanisms^1, 2^. In animal models, these changes recapitulate the hallmarks of human conditions by meeting the criteria of face, construct, and predictive validity^3^.

The hippocampal formation, involved in the activation and feedback of the stress response^4–6^, includes the ventral hippocampus (vHPC), a functionally distinct structure^7, 8^ encoding affective behavior, such as depression^9, 10^ and social avoidance^11^, that mediates the response to antidepressants^12^, and expresses unique patterns of genes^13^. In the vHPC, gene networks have been characterized in animal models each with discrete paradigms of stress, however genomic commonalities in multiple models of stress with distinct construct validity are largely uncharacterized.

Whole-genome RNA-sequencing (RNA-seq) combined with weighted gene coexpression network analysis (WGCNA)^14^ was used to examine transcriptomic signatures across animal models of stress. CORT in drinking water was used to induce susceptibility to stress in wild-type male mice^15^ and heterozygous brain-derived neurotropic factor Val66Met (BDNF het-Met) male mice, which carry a genetic variant that also increases susceptibility to stress^16, 17^. In a separate model, chronic social defeat stress (CSDS) in adulthood, a naturalistic chronic stress model that reflects uncontrollable social aggression^18^, was used to distinguish a behaviorally resilient (RES) and a susceptible (SUS) population of male mice^19^. Mice subjected to CSDS were raised in either standard housing (SH) or enriched environment (EE), which introduced an additional environmental component to stress susceptibility^20, 21^.

Here, the pharmacological, genetic, and environmental effects on behavior and genomics were isolated in either model and then combined using WGCNA, which revealed two networks of converging genes in groups exhibiting either behavioral susceptibility or behavioral resilience.

## METHODS

### Animals

Mice heterozygous for the BDNF allele (het-Met) were generated in the Lee laboratory, as previously described^16^. C57/BL6 mice (WT) and BDNF het-Met mice were obtained by performing in-house breeding. A separate cohort of mice at PND 21 were randomly assigned housing in either SH or EE conditions for 8 weeks. At two months of age WT or BDNF het-Met mice were randomly assigned to either vehicle- (1% ethanol) or CORT- (25mg/l, 1% ethanol) treated groups. At 3-weeks and 4-weeks into the treatment course, mice were tested using the light dark box and splash test, respectively (see Supplementary Materials). After the 8 weeks in their respective conditions, the EE and SH cohort underwent 10 days of CSDS, followed with the social interaction (SI) test (see Supplementary Materials).

### RNA-seq

After the 6-week CORT treatment, mice were killed by cervical dislocation. The vHPC was dissected, immediately flash frozen, and stored at −80 °C. The vHPC is enriched in glucocorticoid receptors, particularly in the dentate gyrus^22^. The ventral dentate gyrus was isolated at P90 from mice that underwent CSDS, the tissues were rapidly removed, flash frozen, and stored at −80 °C. The CORT model utilized three biological replicates per experimental group, with each replicate comprising of RNA pooled from two animals for sequencing by The Rockefeller University Genomics Core Facility. One to seven replicates of RNA sample (not pooled) from each experimental group of animals were randomly selected for the RNA library preparation. When pooling, RNA extracts were combined to reach a final concentration of 100ng/μl. Quality control was performed on the reads obtained from the core and reads with a score of less than 15 were discarded^23, 24^. The reads were then aligned to the GRCm38 genome using the STAR aligner^25^ with Ensembl annotation^26^ and quantified to the gene-level using featureCounts^27^. The read counts were analyzed using the R/Bioconductor framework^28^ (www.R-project.org).

### Sequencing Analysis and Statistics

Behavioral data were analyzed using GraphPad Prism (San Diego, CA, USA) by performing a two-way ANOVA followed by Newman-Keuls *post-hoc* analysis for multiple comparisons. A *p*-value <0.05 was set for statistical significance. Z-score was used to compile complementary variables of behavior (see Supplementary Materials). Differentially expressed gene (DEG) expression analysis used the Limma-Voom package^29^. Overlaps between the differential expression of two independent RNA-seq comparisons were analyzed with a rank-rank hypergeometric overlap (RRHO) analysis^30^. The raw reads were independently processed through WGCNA, a systems biology analysis method for describing the correlation pattern among genes across samples^31^. Enrichment of module DEGs was assessed through Fisher’s exact test corrected for multiple testing (Benjamini-Hochberg False Discovery Rate <0.05) and fold change >1.3. Gene ontology (GO) categories were manually curated from results of the Database for Annotation, Visualization and Integrated Discovery (DAVID) functional annotation cluster tool.

### Data Availability

RNA seq data have been deposited to GEO GSE174664 (https://www.ncbi.nlm.nih.gov/geo/query/acc.cgi?acc=GSE174664) and GSE150812 (https://www.ncbi.nlm.nih.gov/geo/query/acc.cgi?acc=GSE150812). DEGs are included in Supplementary Data Sets. All other data are available upon request.

## RESULTS

### Behavioral phenotyping of male mice under CORT treatment or CSDS combined with housing condition

The cohort of mice maintained on CORT treatment was assessed for anxiety- and depression-like behavior using light-dark box and splash test, respectively. At day 21 of treatment, mice were tested in the light-dark box. CORT induced anxiety-like behavior in WT mice compared to their respective controls when testing the latency to the light box. However, WT mice under CORT showed increased time spent in the light box compared to BDNF het-Met mice (Fig. 1b,c). Mice were maintained on oral CORT, and one week later they were tested using the splash test, where reduced grooming meets predictive validity for increased depression-like behavior^32^. Here, CORT induced depression-like behavior, although this effect was limited to BDNF het-Met mice when testing grooming latency and the number of grooming sessions compared to their controls treated with vehicle (Fig. 1e,g). Thus, although affective behavior was impaired after CORT in both genotypes, WT and BDNF het-Met mice displayed specificity to either test for anxiety-like behavior and depression-like behavior, respectively. To increase the sensitivity and reliability of the behavioral measurements, we applied z-normalization across complementary variables scored in both tests^33^. The distribution of the z-scores showed a cumulative increased anxiety- and depression-like behavior in CORT-treated mice compared to vehicle-treated mice regardless of genotype (Fig.1h).

**Figure 1.**
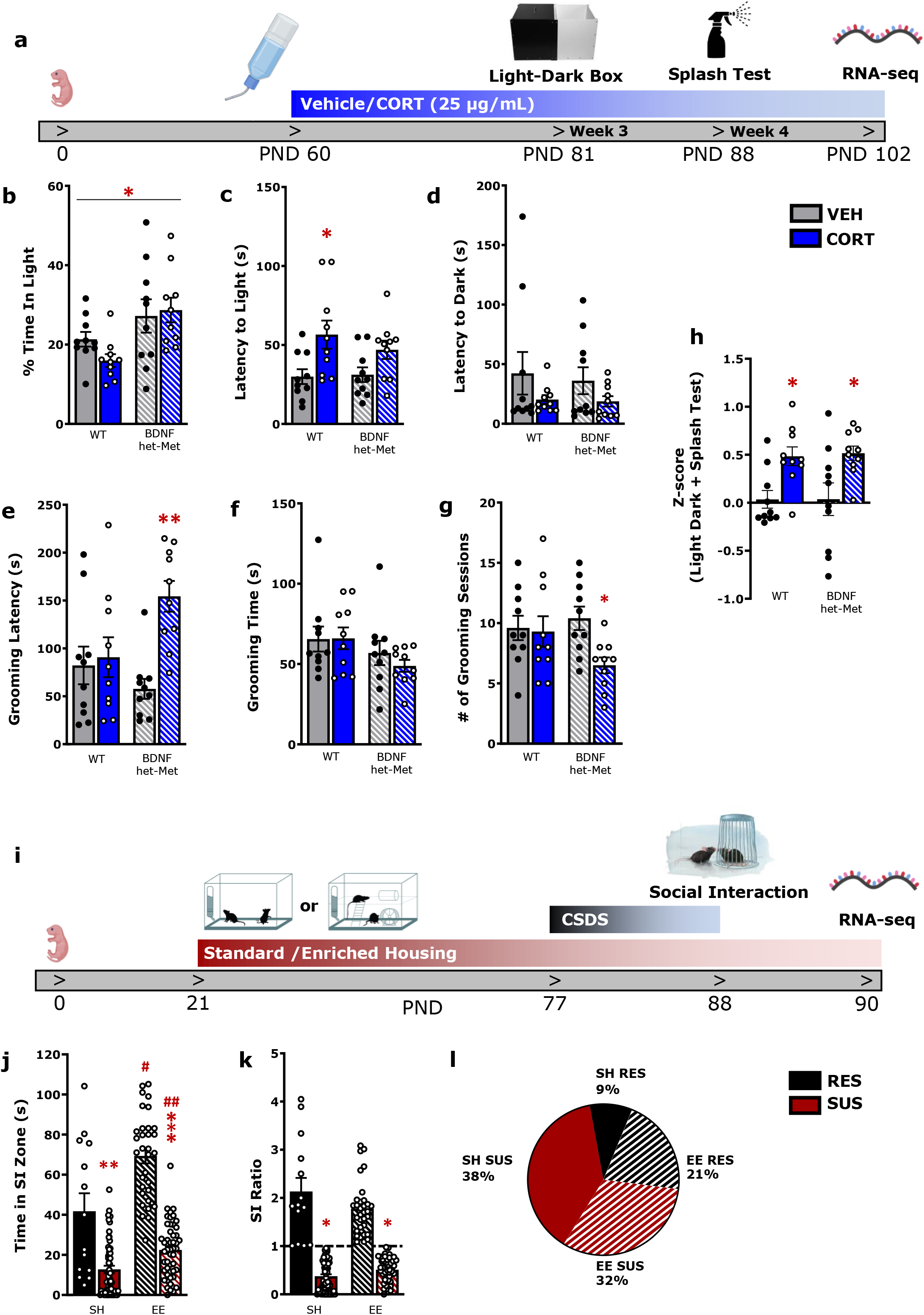
Measurements of emotional behavior in the CORT model and CSDS model show similar face validity in male mice. a) Timeline for CORT treatment experiment. b-d) Light dark test reveals increased anxiety-like behavior in mice treated with CORT. (2-way ANOVA, % Time in Light: genotype: F(1,36) = 10.36, p<0.01; Latency to Light: treatment F(1,36) = 11.47, p < 0.01). e-g) Splash test reveals increased anxiety-like behavior in BDNF het-Met mice treated with CORT (2-way ANOVA, grooming latency: treatment x genotype: F(1,36) = 6.523, p < 0.05; # of grooming sessions: treatment F(1,36) = 4.39, p < 0.05). h) Complementary variables of behavior for both WT and BDNF het-Met were compiled to calculate a z-score, which show that CORT-treated mice displayed higher emotionality scores compared to vehicle-treated mice regardless of genotype (2-way ANOVA, treatment: F(1,36) = 16.44, p < 0.001). i) Timeline for CSDS treatment experiment. j-k) The social interaction reveals decreased anxiety-like behavior in RES mice. (2-way ANOVA, Time in SI Zone: interaction (susceptibility-by-housing) effect: F(1,150) = 7.068, p<0.01; SI Ratio: susceptibility effect: F(1,150) = 243.2, p < 0.001). l) Pie-chart depicting the proportion of mice raised in SH or EE and ranked RES or SUS after CSDS. The proportion of RES mice raised in EE (21%) was higher than the one of RES mice raised in SH (9%) (p<0.05). Columns represent the mean ± S.E.M. of 9–12 determinations per group (a-h) and mean ± S.E.M. of 9–12 determinations per group. */#p < 0.05, **p < 0.01, ***/##p < 0.001. PND: post-natal day, WT: wildtype, BDNF het-Met/hMet: heterozygous for brain-derived neurotropic factor Val66Met, VEH: vehicle, CORT: corticosterone, CSDS: chronic social defeat stress, SI: social interaction, RES: resilient, SUS: susceptible, SH: standard housing, EE: enriched housing

A separate set of mice housed in either SH or EE underwent CSDS and was then assessed for social behavior using the social interaction (SI) test, as a standard to rank mice as SUS or RES to CSDS. SUS mice exhibited CSDS-induced reduction in time spent in the SI zone and the inside/outside interaction ratio that was lower than the significant threshold (=1) compared to RES mice (Fig. 1j,k). Interestingly, in both RES and SUS mice, EE induced increased time spent in the SI zone compared to mice raised in SH (Fig. 1j). The proportion of mice ranked as either SUS or RES considerably changed when mice were raised in SH or EE. Indeed, of the total cohort of mice, 21% were RES and raised in EE compared to only 9% of the cohort ranking RES after SH (Fig. l).

Altogether, these results indicate that chronic treatment with oral CORT or CSDS after alternate housing condition led to behavioral phenotypes consisting of altered affective or social behavior when tested with different paradigms.

### Converging transcriptional responses of oral CORT and CSDS

We next investigated the genomic correlates of these disparate behavioral phenotypes in CORT-treated mice and mice exposed to CSDS that were housed in either SH or EE and examined whether there were converging transcriptional responses. The behavioral findings favored the genomic focus in the vHPC, whose pharmacological modulation meets face validity for the behavioral paradigms reported in this study^34–39^. Using RNA-seq data we identified transcriptional synchrony in DEGs (p<0.05, FC>1.3) between groups based on treatment in drinking water and susceptibility to CSDS. CORT administration induced 220 DEGs in WT and 569 DEGs in BDNF het-Met mice compared to vehicle, with only 62 common DEGs across genotypes. CSDS induced 273 and 238 DEGs between SUS mice compared to RES mice in animals raised in SH and EE, respectively, sharing only 15 common DEGs across housing conditions (Supplementary Data 1). The number of DEGs across comparisons indicated that CORT and CSDS induced gene expression change in the vHPC as a function of genetics and housing condition.

We then used a stratified RRHO test combined with GO to identify the patterns and weight of overlap of DEGs across models and the biological pathways that corresponded to common DEGs of the highest rank between models. Significant overlaps were identified using the point of highest −log10(p-value) from each quadrant as described in Plaiser et al.^30^ (see Supplementary Methods) with non-relevant quadrants shaded in grey. We identified a robust overlap between genes downregulated in WT mice under CORT compared to vehicle-treated mice and downregulated in SUS mice compared to RES mice both raised in SH (max-log10(p-value)=78). We also observed a significant overlap between upregulated genes in WT mice under CORT compared to vehicle and SUS mice compared to RES mice raised in SH (Fig. 2b). The same patterns of overlap, albeit smaller, were also observed when in the analysis WT mice were replaced with BNDF het-Met mice treated with CORT (max-log10(p-value)=76) (Fig. 2c). We also found a significant overlap in DEGs upregulated in WT mice under vehicle and SUS mice raised in EE, compared to CORT and RES mice, respectively (max-log10(p-value)=90) (Fig. 2d). This same pattern of overlap was observed in BDNF het-Met mice treated with vehicle (max-log10(p-value)=115) (Fig. 2e). DEGs from each overlap were grouped according to their biological functions using GO. We found two main macro-networks of genes that converged in discrete comparisons (Supplementary Table 1). The network induced in SUS mice raised in SH and mice treated with CORT of both genotypes was implicated in the regulation of neurotransmission, cytoskeleton function, and brain vascularization, especially regarding oxygen transport and iron binding (Supplementary Table 1; Supplementary Data 2). Specifically, CORT-treated WT and BDNF het-Met mice shared 165 genes with SUS mice raised in SH (Supplementary Fig. 1a; Supplementary Data 2). In the network that included genes concordant in RES mice raised in SH and vehicle-treated mice of both genotypes, GO terms mainly referred to the immune response, the Wnt/PI3K pathway activation, or coded for growth factors (Supplementary Table 1; Supplementary Data 2). Curiously, about 40% of DEGs with these same functions were also expressed in SUS mice raised in EE and vehicle-treated mice regardless of genotype (Supplementary Data 2). Thus, SUS mice raised in EE expressed a novel set of genes that shifted the overlap away from CORT-treated mice. We identified 87 genes that were common in RES mice raised in SH and SUS mice raised in EE and that also matched with vehicle-treated mice regardless of genotype (Supplementary Fig. 1b; Supplementary Data 2). A different set of genes, also involved in neurotransmission, cytoskeleton function, and brain vascularization was common to BDNF het-Met mice under vehicle and SUS mice raised in SH, suggesting that, as opposed to WT, the BDNF Met variant alone moved the DEGs convergence towards SUS mice raised in SH.

**Figure 2.**
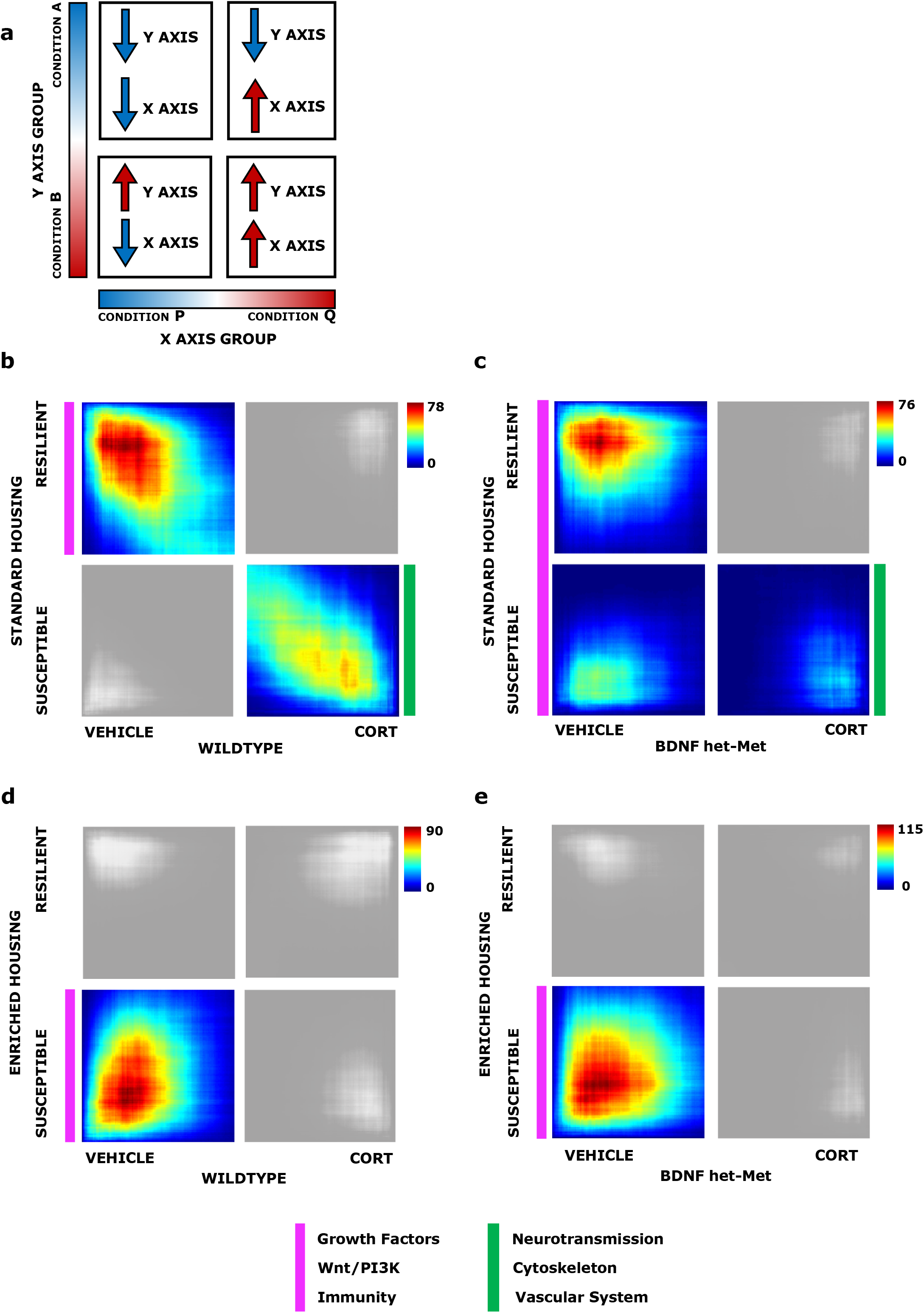
Genomic signatures parallels vulnerability to stress across models and is modulated by housing condition or genotype. a) Representative image of the following threshold free rank-rank hypergeometric overlap comparisons of DEGs. The direction of regulation in each quadrant that corresponds to each variable on each axis is described with an up or down arrow. The upper left quadrant designates co-downregulated genes, lower right quadrant designates co-upregulated genes, and the lower left and upper right quadrants include oppositely regulated genes (up-down and down-up, respectively). Genes along each axis are listed from greatest to least significantly regulated from the outer to middle corners. b-e) Each RRHO contained 17,122 DEGs across the four quadrants. Pixels represent the overlap between the transcriptome of each comparison as noted, with the significance of overlap [-log10(p-value) of a hypergeometric test] color coded. Quadrants that were not significant to the analysis are shaded in grey. b) Comparison of SUS versus RES DEGs in SH with vehicle versus CORT DEGs in WT; RES and vehicle: 738 DEGs; SUS and CORT: 795 DEGs. c) Comparison of SUS versus RES DEGs in SH with vehicle versus CORT DEGs in BDNF het-Met; RES and vehicle: 636 DEGs; SUS and vehicle 522; SUS and CORT: 307 DEGs. d) Comparison of SUS versus RES DEGs in EE with vehicle versus CORT DEGs in WT; SUS and vehicle: 667 DEGs. e) Comparison of SUS versus RES DEGs in EE with vehicle versus CORT DEGs in BDNF het-Met; SUS and vehicle: 870 DEGs. Parallel colored markers indicate macro modules with converging GO terms across quadrants. Genes are listed in Supplementary Table 1.

Together, we showed that CSDS in SUS mice raised in SH induced gene networks common to mice treated with CORT, but EE in SUS mice dramatically reduced this similarity.

### Selection of multimodel gene networks based on co-expression analysis

Moving from the RRHOs predictions, we sought to investigate whether the gene network converging groups met significancy for gene co-expression. Using weighted gene co-expression analysis (WGCNA), we constructed a consensus gene coexpression network and identified 54 gene modules (Supplementary Data 3) that differ in topological overlap, each of which included a unique network of interconnected gene hubs (Fig. 3) based on hierarchal clustering and network preservation (Supplementary Fig. 2)^14^. Each module was assigned an arbitrary color, and we found that the *cyan* and *yellow* module were the only two modules that showed enrichment of DEGs (p<0.05) across the combination of all experimental groups (Fig. 3b). The *cyan* module included upregulated DEGs of mice under CORT of both genotypes and SUS mice raised in SH, compared to vehicle-treated mice and RES mice, respectively. The *yellow* module included DEGs that were upregulated in BDNF Val66Met mice under vehicle and SUS mice, compared to mice under CORT and RES mice, respectively. This pattern of enrichment was consistent with the overlaps observed in the RRHO analysis (Fig. 2b,c,e). Interestingly, groups found in the *cyan* module showed increased stress-related behavior (Fig. 2b,c,h-j), while groups of the *yellow* module exhibited decreased stress-related behavior (Fig. 2c,e,g,i). Together, these analyses suggest that the *cyan* and *yellow* modules are implicated in governing the emergence of stress-related phenotypes in both mice treated with oral CORT and mice raised in different housing conditions prior exposure to CSDS.

**Figure 3.**
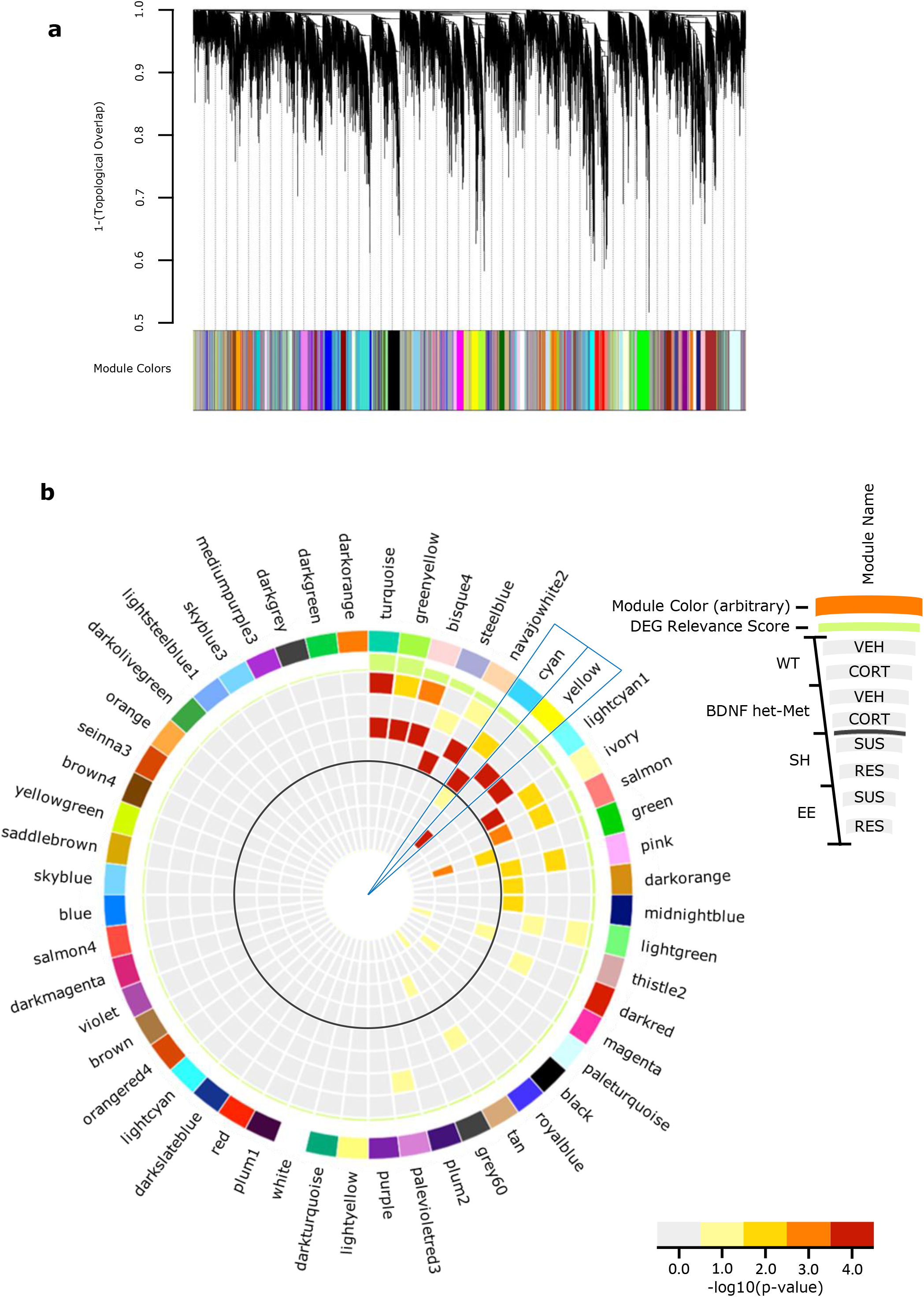
Identification of Shared-Model Coexpression Networks and Key Modules. a) Consensus coexpression network analysis identified 54 coexpressed modules in both the CORT and CSDS experiments via hierarchal gene clustering on TOM-based dissimilarity and branch cutting using the top-down dynamic tree cut method. Each module is assigned a unique color identifier along the bottom of the dendrogram. Dendrograms demonstrate average linkage hierarchical clustering of genes based on the calculated topological overlap values. b) Circos plot showing module name (ring 1), color (ring 2), a differential expression relevance score; calculated from the average enrichment of DEGs across all groups, with increasing bar height indicating increased average total enrichment (ring 3). Bar color indicates the significance for genes upregulated for each condition (vehicle compared to CORT, CORT compared to vehicle, SUS compared to RES, and RES compared to SUS) with warmer colors signifying increasing −log10(p-value). The solid grey circle indicates the separation between the DEGs of the model of oral CORT in the outer shell and the DEGs of the model of differential housing before CSDS in the inner shell. The *cyan* and *yellow* modules are indicated as they have higher DEG average relevance, along with significant enrichment between models. WT VEH up (ring 4), WT CORT up (ring 5), BDNF het-Met VEH up (ring 6), BDNF het-Met CORT up (ring 7), SH SUS up (ring 8), SH RES up (ring 9), EE SUS up (ring 10), EE RES up (ring 11). VEH: vehicle, CORT: corticosterone, RES: resilient, SUS: susceptible, SH: standard housing, EE: enriched housing.

### Probing the intramodular network structure and function of key modules

To identify specific drivers particular to our target modules, we reconstructed the network structure of genes within each module based on their coexpression-based interconnectivity and then identified ‘hub genes’ (Fig 4a,b). Hub genes (or key drivers) are highly connected genes within a module that are expected to control the expression of many other module members, though this prediction is derived from non-directed correlational analysis. The three highest connected hub genes within the *cyan* module included *Wdr7, Camsap2*, and *Dnajc6* (Supplementary Data 4). Other genes are included with the hub analysis and while lower in connectivity are DEGs and shared across groups: *Spink8* and *Krt73* are upregulated in both WT and BDNF het-Met mice administered CORT, *Alox12b* is upregulated in both WT mice administered CORT and SUS mice raised in SH, and *Adamts14* is upregulated in both BDNF het-Met mice administered CORT and SUS mice raised in SH (Supplementary Data 5). The three highest connected hub genes within the *yellow* module included *Arap2, Vwc2*, and *Cpne9* (Supplementary Data 4). Additional hubs within the top 10 highest connected genes included *Plcβ4*, which is a DEG upregulated in SUS mice raised in EH, and *Tnnt1* and *Chn2*, which are DEGs upregulated in BDNF het-Met mice administered CORT. *Usp43*, while relatively lower in connectivity, was differentially expressed in both target groups of the *yellow* module (Supplementary Data 5).

**Figure 4.**
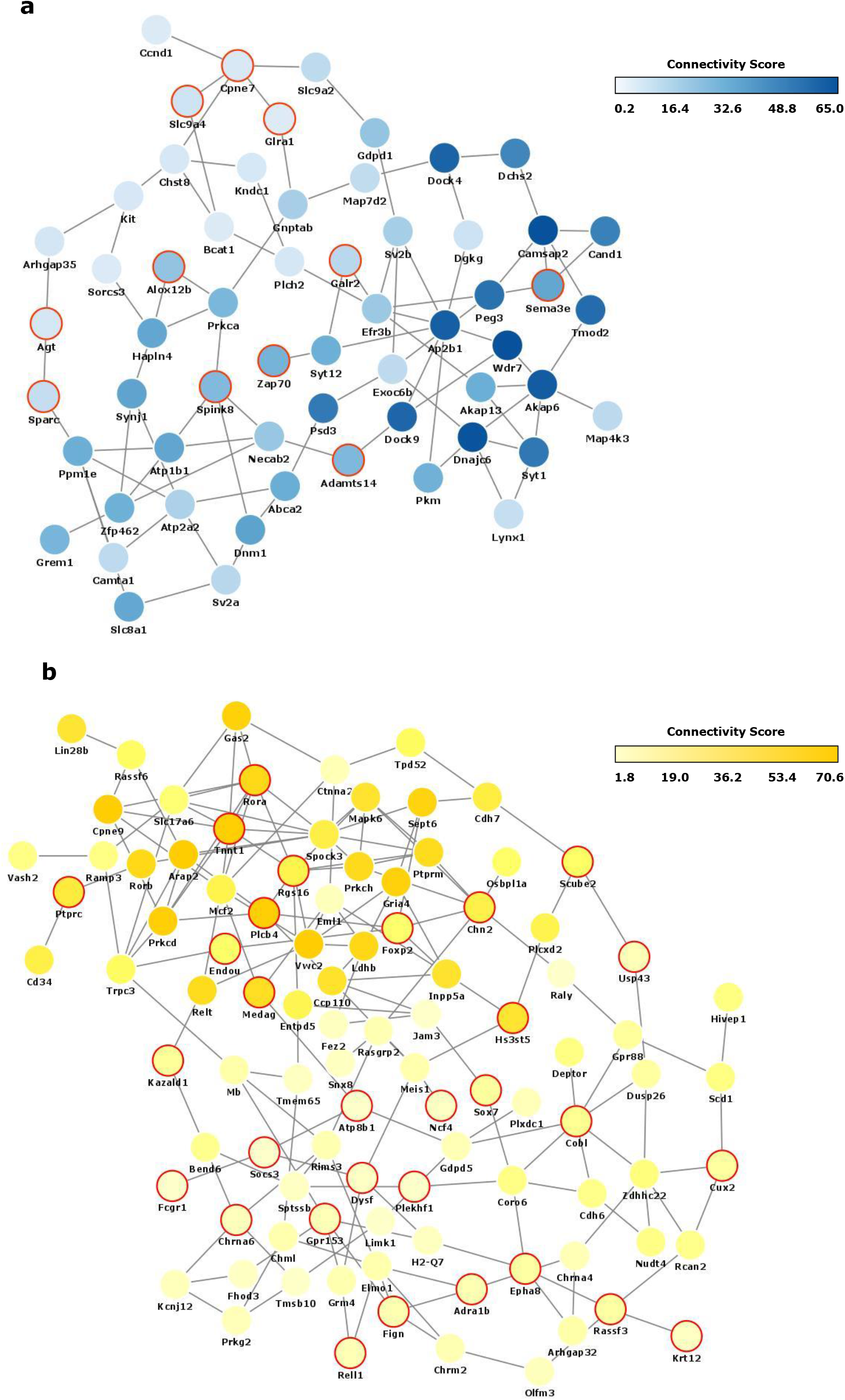
Co-expression Networks of *Yellow* and *Cyan* Modules. a) Network plot of hub genes within the *cyan* module. b) Network plot of hub genes within the *yellow* module. Most significant nodes were selected from the most connected genes greater than 1 standard deviation from the average (μ + σ) when organized by decreasing KME score. Node color is proportional to the node’s level of connectivity within the hub. Red outline indicates a differentially expressed gene.

## DISCUSSION

We investigated functional gene networks in animal models of stress with different construct validity by conducting coexpression analysis of DEGs. Unique hub genes and coexpression networks were exclusively found in mice exhibiting either behavioral susceptibility or resilience to stress.

CORT induced increased affective behavior in both WT and BDNF het-Met male mice, indicating that both genotypes showed behavioral susceptibility to CORT as previously reported^40–42^. This finding was reinforced by compiling multiple behavioral variables that meet converging face validity^43, 44^, a translational application that recapitulates illness definition (i.e. a syndrome as a collection of variable symptoms), which incorporates converging variable of symptoms especially in affective disorders^33^. In this study, behavioral susceptibility to CSDS^45^ varied significantly whether mice were raised in SH or EE and was consistent with findings showing that EE reduces emotional lability^46^ even when experienced prior to stress^47^. SI analyzes a unique trait of depression-like behavior and shows pharmacological validity for stress-related disorders^48^.

To determine whether there existed converging genetic patterns that drove these disparate stress-related phenotypes, we focus on the vHPC, a brain region whose pharmacological and anatomical manipulation has been shown to induce effects in the same behavioral tests utilized in this study^34–38^. Indeed, CORT treatment and environmental enrichment have been associated with selective changes in neurogenesis^49^ and gene expression^6^ in the vHPC. Others have reported remarkable stress-induced gene expression differences across multiple brain regions, such as the nucleus accumbens or the medial prefrontal cortex, especially after CSDS^4, 50, 51^. So, it remains to investigate whether other brain regions would also show converging genomic signatures across the paradigms reported here.

By referencing novel RNA-seq data from the vHPC, we used RRHO combined with GO to depict the multimodel trends of transcriptional regulation and biological function. One macro network, which included genes that participates in the immune response, Wnt/PI3K pathway activation, or coding for growth factors was found in quadrants associated with resilience, while a distinct macro network implicated in neurotransmission, cytoskeleton function, and brain vascularization, was expressed in quadrants associated with susceptibility. The immune system is influenced by stress and glucocorticoids^52, 53^, and along with Wnt signaling and growth factors has been implicated in depression pathogenesis^52–54^. Interestingly, disruption of cytoskeleton functions^55^ and alteration of brain vascularization and vessels^56^, have also been associated with neuroanatomical changes of the hippocampus, depression-like behavior, and susceptibility to neuropsychiatric disorder.

The lack in common biological pathways affected in the RRHO overlaps of interest is consistent with the distinct genomic landscape observed in stress susceptibility and resilience^4, 6^. We observed a smaller, compared to WT mice, but unique overlap between SUS mice and BDNF het-Met mice treated with CORT, suggesting that exclusive sets of genes are induced by stress in BDNF het-Met mice, as previously demonstrated^57, 58^. The most significant overlap was between SUS mice raised in EE and mice under vehicle of both genotypes. Thus, EE in SUS mice profoundly reduced the similarity between CORT-treated and SUS mice, shifting the overlap instead to vehicle-treated mice, indicating that EE reverses those genomic signatures of stress typically associated with susceptibility. This evidence complements findings that EE promotes hippocampal neurogenesis and restores behavior after social defeat^59^.

While this analysis elucidates individual DEGs, we then used differential connectivity to examine multi-dimensional alterations between pairs of genes, as susceptibility to stress-related disorders is characterized by fundamental changes in the architecture of transcriptional networks^60^. Accordingly, WGCNA analysis showed that two modules, *yellow* and *cyan*, were enriched in DEGs across experimental groups. This network-based approach allowed us to identify several hub genes, which according to their definition are more likely to drive the function of the entire network^14, 61^. The significant enrichment of the *cyan* module indicated that CORT treatment regardless of genotype exhibited gene expression synchrony with SUS mice raised in SH prior CSDS, an expression pattern that reflected increased affective behavior observed selectively in these groups. The most highly connected hub gene of the *cyan* module was *Wdr7*, whose altered expression was found in subjects with depression and alcohol dependence comorbidity^62^. The intramodular network of *Wdr7* included *Adamts14* and *Dock4*, two markers of neuropsychiatric disorder susceptibility^63, 64^. The *yellow* module demonstrated genomic synchrony in mice exhibiting low affective behavior score, namely BDNF het-Met mice under vehicle and SUS mice raised in EE, emphasizing previous evidence showing that BDNF het-Met mice express genes associated with stress coping even in the absence of applied stressors^58^. The top hub regulator of the *yellow* module, *Arap2*, encodes for pro-survival functions, such as Akt activity^65^, glucose uptake, and sphingolipid metabolism^66^, and whose dysregulation leads to impaired affective behaviors in animal models of depression^67, 68^. In the *yellow* module, *Arap2* was interconnected with other resilience-associated genes. For example, increased expression of *Tnnt1* was demonstrated in mice after injection with ketamine^69^, an NMDA receptor antagonist used against treatment-resistant depression^70^. The intramodular network of *Arap2* also included *Plcβ4* and *Rora*, whose functional expression prevents anxiety^71^ or major depression^72^. Therefore, the *cyan* and *yellow* modules included hubs that associate with stress vulnerability or survival, respectively. We offer a model that summarizes converging intramodular networks that are regulated in those behavioral phenotypes that also converge for increased or decreased affective behavior. More properly, mice that express the *cyan* module display behavioral susceptibility, while mice that express the *yellow* module exhibit behavioral resilience (Fig. 5).

**Figure 5.**
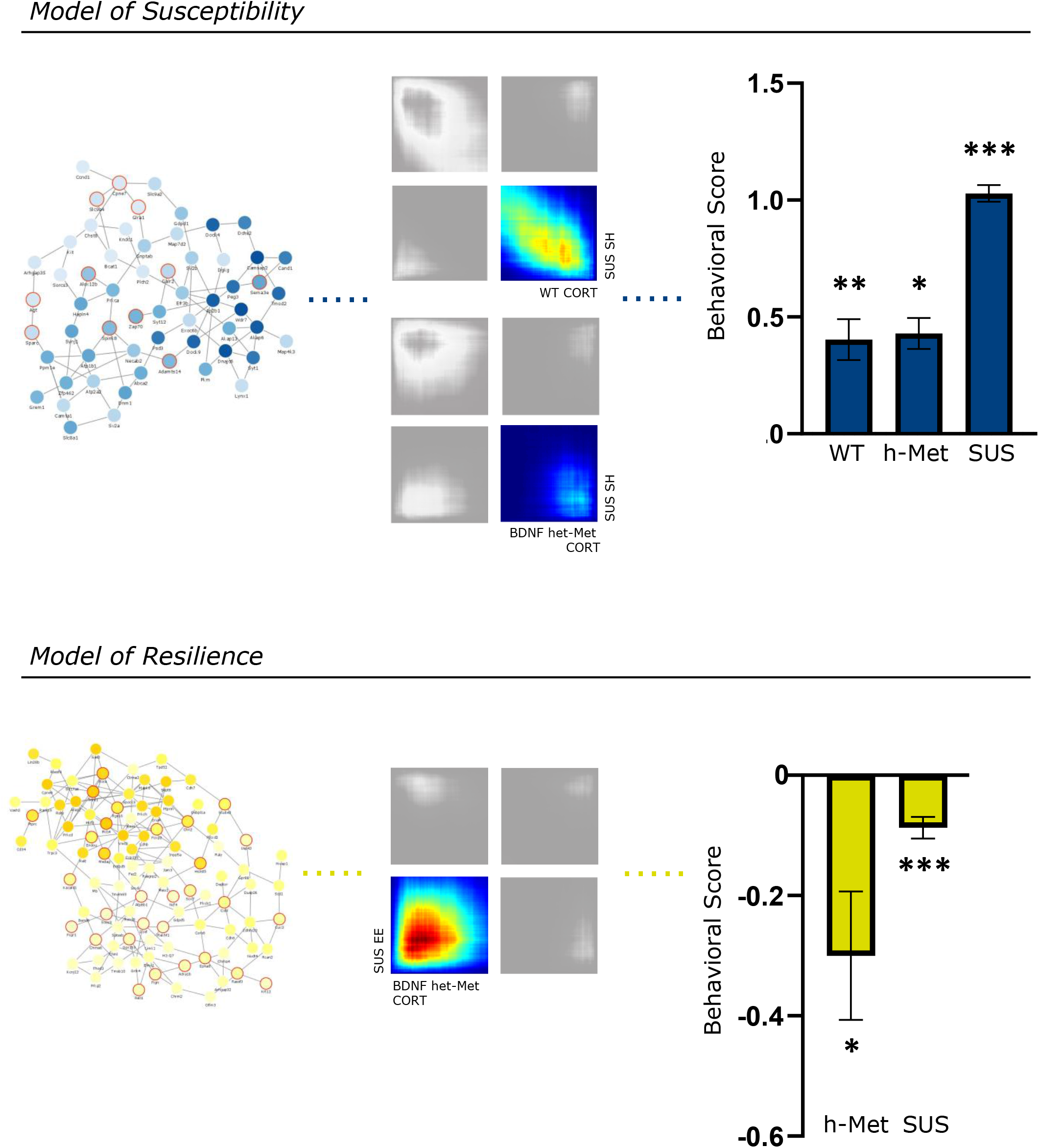
Summary of Transcriptomic and Behavioral Syncrony. Enrichment in hub genes and DEGs in the *cyan* module, matching RRHOs, and behavioral susceptibility to stress are found in CORT-treated mice of both genotypes and SUS mice raised in SH. Enrichment in hub genes and DEGs in the *yellow* module, concordance in RRHO gene expression, and behavioral resilience to stress (*yellow* column) are found in BDNF het-Met treated with vehicle and SUS mice raised in EE. The behavioral score for each group was normalized to its respective control: WT CORT animals (vs. WT vehicle, t=3.356, df=17.95, p<0.01), BDNF het-Met mice treated with vehicle for BDNF het-Met CORT animals (t=2.579, df=12.30, p<0.05), RES in SH for SH SUS mice (t=5.454, df=13.98, p<0.001), BDNF het-Met CORT for BDNF het-Met vehicle animals (t=2.579, df=12.30, p<0.05), and SH SUS for EE SUS animals (t=3.455, df=104.6, p<0.001). Behavioral score for each animal is calculated as: 1-z/ā, where 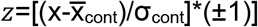 and x=percentage time in the light, latency to dark, latency to light, grooming time, number of grooming sessions, grooming latency, time in social interaction zone, or SI ratio, and *α=z* for each respective control mouse. After calculation, *α* and *z* values where normalized to the lowest *z* score. Columns represent the mean ± S.E.M. of 9–12 determinations per group. *p < 0.05, **p < 0.01, ***p < 0.001. DEG: differentially expressed gene, WT: wildtype, BDNF het-Met/hMet: heterozygous for brain-derived neurotropic factor Val66Met, CORT: corticosterone, SUS: susceptible, SH: standard housing, EE: enriched housing.

Here, we utilized multiple variables contributing to stress-related disorders, such as genetic inheritance and environment. When comparing pharmacological and environmental models, one should consider unique physiological mechanisms due to disparate stress-inducing conditions. Nevertheless, some genetic pathways may act as an independent core or even modulate the multifactorial physiological responses downstream, making it possible to genomically funnel the diversity of the models used to study stress-related disorders. While this study focuses on males, data previously reported in females, from the same cohort of males included here, show that CORT reduces, rather than increases, affective behavior selectively in BDNF het-Met mice and induces gene expression patterns that profoundly differ from those described in males^73^. Therefore, with the intent to exclusively combine models with converging face validity for susceptibility, females were not included after observing a behavioral phenotype that opposed all the groups included in this study. These sex differences will also have to focus on understanding genomic signatures underlying disparate face validity rather than converging behavioral and molecular traits. Furthermore, data in females exposed to CSDS and alternate housing are scarce, and the comparison with females treated with CORT warrants a separate investigation. However, one should also note that CSDS in females requires a behavioral paradigm distinct from the one used in males^74^, introducing an additional variable of comparison. Without neglecting the importance of mosaic gene networks in complex illnesses, further studies may also secure the identification of targets at the single gene level, using silencing/activation methods to dissect the gene driver(s) of the modular network that corroborate a causal link with the behavioral phenotypes that we reported.

Stress-related disorders are multifaceted, constraining our understanding of their molecular etiologies^75^. Investigating the holistic network of the genome associated with stress-related disorders could shift focus to key biological systems and facilitate the identification of novel therapeutic targets, with the scope of constructing interventions to target whole connectivity networks rather than individual DEGs^60^. The distinction between the effect of different types of stress on animal behavior and physiology remains largely unexplored but it has practical implications in order to recapitulate the complexity of human conditions in preclinical models^3^. This supports the advancement of multimodel metanalysis for the dissection of behavioral circuits and their molecular correlates as targets for novel therapies in psychiatric disorders.

## ACKNOWLEDGEMENTS

This work was supported by the Hope for Depression Research Foundation (H.A., B.S.M., and M.J.M.). We thank the Rockefeller University Genomics Core Facility and the Rockefeller University Bioinformatics Resource Center for performing the RNA-sequencing and for computing and analyzing the datasets, respectively. In memory of Bruce S. McEwen.

## COMPETING INTERESTS

The authors declare no competing interests.

## Notes

https://www.ncbi.nlm.nih.gov/geo/query/acc.cgi?acc=GSE174664

https://www.ncbi.nlm.nih.gov/geo/query/acc.cgi?acc=GSE150812

### Competing Interest Statement

The authors have declared no competing interest.

## REFERENCES

1. Gray JD, Kogan JF, Marrocco J, McEwen BS. Genomic and epigenomic mechanisms of glucocorticoids in the brain. Nat Rev Endocrinol. 2017;13(11):661–73.

2. McEwen BS, Bowles NP, Gray JD, Hill MN, Hunter RG, Karatsoreos IN, et al. Mechanisms of stress in the brain. Nat Neurosci. 2015;18(10):1353–63.

3. Nestler EJ, Hyman SE. Animal models of neuropsychiatric disorders. Nat Neurosci. 2010;13(10):1161–9.

4. Bagot RC, Parise EM, Peña CJ, Zhang HX, Maze I, Chaudhury D, et al. Ventral hippocampal afferents to the nucleus accumbens regulate susceptibility to depression. Nat Commun. 2015;6:7062.

5. McEwen BS. Stress, sex, and neural adaptation to a changing environment: mechanisms of neuronal remodeling. Ann N Y Acad Sci. 2010;1204 Suppl(Suppl):E38–59.

6. Zhang TY, Keown CL, Wen X, Li J, Vousden DA, Anacker C, et al. Environmental enrichment increases transcriptional and epigenetic differentiation between mouse dorsal and ventral dentate gyrus. Nat Commun. 2018;9(1):298.

7. Fanselow MS, Dong HW. Are the dorsal and ventral hippocampus functionally distinct structures? Neuron. 2010;65(1):7–19.

8. Kjelstrup KG, Tuvnes FA, Steffenach H-A, Murison R, Moser EI, Moser M-B. Reduced Fear Expression after Lesions of the Ventral Hippocampus. Proceedings of the National Academy of Sciences of the United States of America. 2002;99(16):10825–30.

9. Jacinto LR, Reis JS, Dias NS, Cerqueira JJ, Correia JH, Sousa N. Stress affects theta activity in limbic networks and impairs novelty-induced exploration and familiarization. Front Behav Neurosci. 2013;7:127.

10. Nalloor R, Bunting KM, Vazdarjanova A. Altered hippocampal function before emotional trauma in rats susceptible to PTSD-like behaviors. Neurobiol Learn Mem. 2014;112:158–67.

11. Markham CM, Taylor SL, Huhman KL. Role of amygdala and hippocampus in the neural circuit subserving conditioned defeat in Syrian hamsters. Learn Mem. 2010;17(2):109–16.

12. Kheirbek MA, Hen R. Dorsal vs ventral hippocampal neurogenesis: implications for cognition and mood. Neuropsychopharmacology. 2011;36(1):373–4.

13. Cembrowski MS, Wang L, Sugino K, Shields BC, Spruston N. Hipposeq: a comprehensive RNA-seq database of gene expression in hippocampal principal neurons. Elife. 2016;5:e14997.

14. Zhang B, Horvath S. A general framework for weighted gene co-expression network analysis. Stat Appl Genet Mol Biol. 2005;4:Article17.

15. Karatsoreos IN, Bhagat SM, Bowles NP, Weil ZM, Pfaff DW, McEwen BS. Endocrine and physiological changes in response to chronic corticosterone: a potential model of the metabolic syndrome in mouse. Endocrinology. 2010;151(5):2117–27.

16. Chen ZY, Jing D, Bath KG, Ieraci A, Khan T, Siao CJ, et al. Genetic variant BDNF (Val66Met) polymorphism alters anxiety-related behavior. Science. 2006;314(5796):140–3.

17. Notaras M, van den Buuse M. Neurobiology of BDNF in fear memory, sensitivity to stress, and stress-related disorders. Mol Psychiatry. 2020;25(10):2251–74.

18. Kollack-Walker S, Don C, Watson SJ, Akil H. Differential expression of c-fos mRNA within neurocircuits of male hamsters exposed to acute or chronic defeat. J Neuroendocrinol. 1999;11(7):547–59.

19. Krishnan V, Han MH, Graham DL, Berton O, Renthal W, Russo SJ, et al. Molecular adaptations underlying susceptibility and resistance to social defeat in brain reward regions. Cell. 2007;131(2):391–404.

20. Laviola G, Hannan AJ, Macrì S, Solinas M, Jaber M. Effects of enriched environment on animal models of neurodegenerative diseases and psychiatric disorders. Neurobiol Dis. 2008;31(2):159–68.

21. Schloesser RJ, Lehmann M, Martinowich K, Manji HK, Herkenham M. Environmental enrichment requires adult neurogenesis to facilitate the recovery from psychosocial stress. Mol Psychiatry. 2010;15(12):1152–63.

22. Romeo RD, Ali FS, Karatsoreos IN, Bellani R, Chhua N, Vernov M, et al. Glucocorticoid Receptor mRNA Expression in the Hippocampal Formation of Male Rats before and after Pubertal Development in Response to Acute or Repeated Stress. Neuroendocrinology. 2008;87(3):160–7.

23. Chen S, Zhou Y, Chen Y, Gu J. fastp: an ultra-fast all-in-one FASTQ preprocessor. Bioinformatics. 2018;34(17):i884–i90.

24. Ewels P, Magnusson M, Lundin S, Käller M. MultiQC: summarize analysis results for multiple tools and samples in a single report. Bioinformatics. 2016;32(19):3047–8.

25. Dobin A, Davis CA, Schlesinger F, Drenkow J, Zaleski C, Jha S, et al. STAR: ultrafast universal RNA-seq aligner. Bioinformatics. 2013;29(1):15–21.

26. Cunningham F, Achuthan P, Akanni W, Allen J, Amode M R, Armean IM, et al. Ensembl 2019. Nucleic Acids Research. 2018;47(D1):D745–D51.

27. Liao Y, Smyth GK, Shi W. The Subread aligner: fast, accurate and scalable read mapping by seed-and-vote. Nucleic Acids Res. 2013;41(10):e108.

28. Huber W, Carey VJ, Gentleman R, Anders S, Carlson M, Carvalho BS, et al. Orchestrating high-throughput genomic analysis with Bioconductor. Nature Methods. 2015;12(2):115–21.

29. Ritchie ME, Phipson B, Wu D, Hu Y, Law CW, Shi W, et al. limma powers differential expression analyses for RNA-sequencing and microarray studies. Nucleic Acids Research. 2015;43(7):e47–e.

30. Plaisier SB, Taschereau R, Wong JA, Graeber TG. Rank-rank hypergeometric overlap: identification of statistically significant overlap between gene-expression signatures. Nucleic Acids Res. 2010;38(17):e169.

31. Langfelder P, Horvath S. WGCNA: an R package for weighted correlation network analysis. BMC Bioinformatics. 2008;9:559.

32. Isingrini E, Camus V, Le Guisquet AM, Pingaud M, Devers S, Belzung C. Association between repeated unpredictable chronic mild stress (UCMS) procedures with a high fat diet: a model of fluoxetine resistance in mice. PLoS One. 2010;5(4):e10404.

33. Guilloux JP, Seney M, Edgar N, Sibille E. Integrated behavioral z-scoring increases the sensitivity and reliability of behavioral phenotyping in mice: relevance to emotionality and sex. J Neurosci Methods. 2011;197(1):21–31.

34. Bannerman DM, Grubb M, Deacon RMJ, Yee BK, Feldon J, Rawlins JNP. Ventral hippocampal lesions affect anxiety but not spatial learning. Behavioural Brain Research. 2003;139(1):197–213.

35. McHugh SB, Deacon RMJ, Rawlins JNP, Bannerman DM. Amygdala and Ventral Hippocampus Contribute Differentially to Mechanisms of Fear and Anxiety. Behavioral Neuroscience. 2004;118(1):63–78.

36. Marrocco J, Mairesse J, Ngomba RT, Silletti V, Van Camp G, Bouwalerh H, et al. Anxiety-Like Behavior of Prenatally Stressed Rats Is Associated with a Selective Reduction of Glutamate Release in the Ventral Hippocampus. The Journal of Neuroscience. 2012;32(48):17143–54.

37. Marrocco J, Reynaert M-L, Gatta E, Gabriel C, Mocaër E, Di Prisco S, et al. The Effects of Antidepressant Treatment in Prenatally Stressed Rats Support the Glutamatergic Hypothesis of Stress-Related Disorders. The Journal of Neuroscience. 2014;34(6):2015–24.

38. Sams-Dodd F, Lipska BK, Weinberger DR. Neonatal lesions of the rat ventral hippocampus result in hyperlocomotion and deficits in social behaviour in adulthood. Psychopharmacology. 1997;132(3):303–10.

39. Anacker C, Luna VM, Stevens GS, Millette A, Shores R, Jimenez JC, et al. Hippocampal neurogenesis confers stress resilience by inhibiting the ventral dentate gyrus. Nature. 2018;559(7712):98–102.

40. Ding H, Cui XY, Cui SY, Ye H, Hu X, Zhao HL, et al. Depression-like behaviors induced by chronic corticosterone exposure via drinking water: Time-course analysis. Neurosci Lett. 2018;687:202–6.

41. Kinlein SA, Wilson CD, Karatsoreos IN. Dysregulated hypothalamic-pituitary-adrenal axis function contributes to altered endocrine and neurobehavioral responses to acute stress. Front Psychiatry. 2015;6:31.

42. Klug M, Hill RA, Choy KH, Kyrios M, Hannan AJ, van den Buuse M. Long-term behavioral and NMDA receptor effects of young-adult corticosterone treatment in BDNF heterozygous mice. Neurobiol Dis. 2012;46(3):722–31.

43. Crawley J, Goodwin FK. Preliminary report of a simple animal behavior model for the anxiolytic effects of benzodiazepines. Pharmacol Biochem Behav. 1980;13(2):167–70.

44. Yalcin I, Aksu F, Belzung C. Effects of desipramine and tramadol in a chronic mild stress model in mice are altered by yohimbine but not by pindolol. European Journal of Pharmacology. 2005;514(2):165–74.

45. Nestler EJ, Gould E, Manji H, Buncan M, Duman RS, Greshenfeld HK, et al. Preclinical models: status of basic research in depression. Biol Psychiatry. 2002;52(6):503–28.

46. Kappeler L, Meaney MJ. Enriching stress research. Cell. 2010;142(1):15–7.

47. Cymerblit-Sabba A, Lasri T, Gruper M, Aga-Mizrachi S, Zubedat S, Avital A. Prenatal Enriched Environment improves emotional and attentional reactivity to adulthood stress. Behav Brain Res. 2013;241:185–90.

48. Berton O, McClung CA, Dileone RJ, Krishnan V, Renthal W, Russo SJ, et al. Essential role of BDNF in the mesolimbic dopamine pathway in social defeat stress. Science. 2006;311(5762):864–8.

49. Workman JL, Chan MYT, Galea LAM. Prior high corticosterone exposure reduces activation of immature neurons in the ventral hippocampus in response to spatial and nonspatial memory. Hippocampus. 2015;25(3):329–44.

50. Labonté B, Engmann O, Purushothaman I, Menard C, Wang J, Tan C, et al. Sex-specific transcriptional signatures in human depression. Nat Med. 2017;23(9):1102–11.

51. Vialou V, Bagot RC, Cahill ME, Ferguson D, Robison AJ, Dietz DM, et al. Prefrontal cortical circuit for depression- and anxiety-related behaviors mediated by cholecystokinin: role of ΔFosB. J Neurosci. 2014;34(11):3878–87.

52. Miller AH, Raison CL. The role of inflammation in depression: from evolutionary imperative to modern treatment target. Nat Rev Immunol. 2016;16(1):22–34.

53. Horowitz MA, Cattaneo A, Cattane N, Lopizzo N, Tojo L, Bakunina N, et al. Glucocorticoids prime the inflammatory response of human hippocampal cells through up-regulation of inflammatory pathways. Brain Behav Immun. 2020;87:777–94.

54. Voleti B, Duman RS. The roles of neurotrophic factor and Wnt signaling in depression. Clin Pharmacol Ther. 2012;91(2):333–8.

55. Molteni R, Calabrese F, Chourbaji S, Brandwein C, Racagni G, Gass P, et al. Depression-prone mice with reduced glucocorticoid receptor expression display an altered stress-dependent regulation of brain-derived neurotrophic factor and activity-regulated cytoskeleton-associated protein. J Psychopharmacol. 2010;24(4):595–603.

56. Baruah J, Vasudevan A. The Vessels Shaping Mental Health or Illness. Open Neurol J. 2019;13:1–9.

57. Gray JD, Rubin TG, Kogan JF, Marrocco J, Weidmann J, Lindkvist S, et al. Translational profiling of stress-induced neuroplasticity in the CA3 pyramidal neurons of BDNF Val66Met mice. Mol Psychiatry. 2018;23(4):904–13.

58. Marrocco J, Petty GH, Ríos MB, Gray JD, Kogan JF, Waters EM, et al. A sexually dimorphic pre-stressed translational signature in CA3 pyramidal neurons of BDNF Val66Met mice. Nat Commun. 2017;8(1):808.

59. Lehmann ML, Brachman RA, Martinowich K, Schloesser RJ, Herkenham M. Glucocorticoids orchestrate divergent effects on mood through adult neurogenesis. J Neurosci. 2013;33(7):2961–72.

60. Bagot RC, Cates HM, Purushothaman I, Lorsch ZS, Walker DM, Wang J, et al. Circuit-wide Transcriptional Profiling Reveals Brain Region-Specific Gene Networks Regulating Depression Susceptibility. Neuron. 2016;90(5):969–83.

61. Ivliev AE, t Hoen PA, Sergeeva MG. Coexpression network analysis identifies transcriptional modules related to proastrocytic differentiation and sprouty signaling in glioma. Cancer Res. 2010;70(24):10060–70.

62. Edwards AC, Aliev F, Bierut LJ, Bucholz KK, Edenberg H, Hesselbrock V, et al. Genome-wide association study of comorbid depressive syndrome and alcohol dependence. Psychiatr Genet. 2012;22(1):31–41.

63. Alkelai A, Lupoli S, Greenbaum L, Kohn Y, Kanyas-Sarner K, Ben-Asher E, et al. DOCK4 and CEACAM21 as novel schizophrenia candidate genes in the Jewish population. Int J Neuropsychopharmacol. 2012;15(4):459–69.

64. Galfalvy H, Haghighi F, Hodgkinson C, Goldman D, Oquendo MA, Burke A, et al. A genome-wide association study of suicidal behavior. Am J Med Genet B Neuropsychiatr Genet. 2015;168(7):557–63.

65. Luo R, Chen PW, Kuo JC, Jenkins L, Jian X, Waterman CM, et al. ARAP2 inhibits Akt independently of its effects on focal adhesions. Biol Cell. 2018;110(12):257–70.

66. Chaudhari A, Håversen L, Mobini R, Andersson L, Ståhlman M, Lu E, et al. ARAP2 promotes GLUT1-mediated basal glucose uptake through regulation of sphingolipid metabolism. Biochim Biophys Acta. 2016;1861(11):1643–51.

67. Detka J, Kurek A, Basta-Kaim A, Kubera M, Lasoń W, Budziszewska B. Elevated brain glucose and glycogen concentrations in an animal model of depression. Neuroendocrinology. 2014;100(2-3):178–90.

68. Lin CH, Yeh SH, Leu TH, Chang WC, Wang ST, Gean PW. Identification of calcineurin as a key signal in the extinction of fear memory. J Neurosci. 2003;23(5):1574–9.

69. Lowe X, Wyrobek A. Characterization of the early CNS stress biomarkers and profiles associated with neuropsychiatric diseases. Curr Genomics. 2012;13(6):489–97.

70. Krystal JH, Charney DS, Duman RS. A New Rapid-Acting Antidepressant. Cell. 2020;181(1):7.

71. Shin J, Gireesh G, Kim SW, Kim DS, Lee S, Kim YS, et al. Phospholipase C beta 4 in the medial septum controls cholinergic theta oscillations and anxiety behaviors. J Neurosci. 2009;29(49):15375–85.

72. Terracciano A, Tanaka T, Sutin AR, Sanna S, Deiana B, Lai S, et al. Genome-wide association scan of trait depression. Biol Psychiatry. 2010;68(9):811–7.

73. Marrocco J, Einhorn NR, Caradonna SG, Le Floch C, Lihagen A, Petty GH, et al., editors. Transcriptional profiling of the stressed hippocampus: Does sex make a difference? Society for Neurosceince; 2019 October 23, 2019; Chicago, IL: 2019 Neurosceince Meeting Planner.

74. Harris AZ, Atsak P, Bretton ZH, Holt ES, Alam R, Morton MP, et al. A Novel Method for Chronic Social Defeat Stress in Female Mice. Neuropsychopharmacology. 2018;43(6):1276–83.

75. Rush AJ. The varied clinical presentations of major depressive disorder. J Clin Psychiatry. 2007;68 Suppl 8:4–10.

