## Supplementary material for "Genomic Modules and Intramodular Network Convergency of Susceptibility and Resilience in Multimodeled Stress in Male Mice": Marrocco_Supplementary_Figures.pdf

**a**

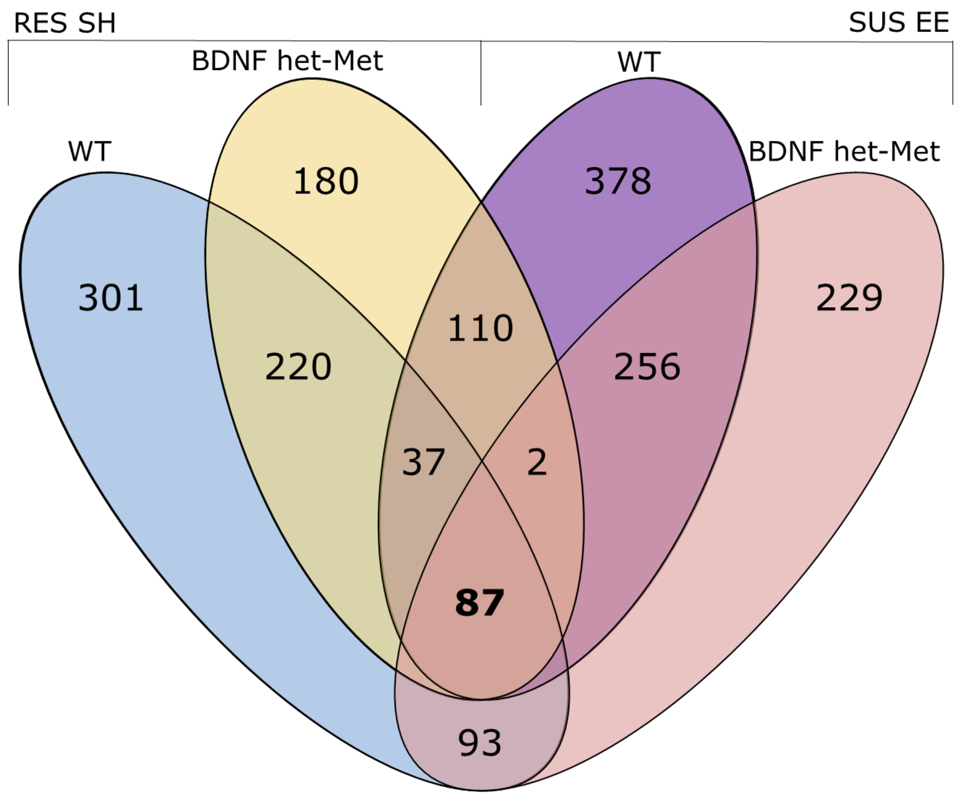


**b**

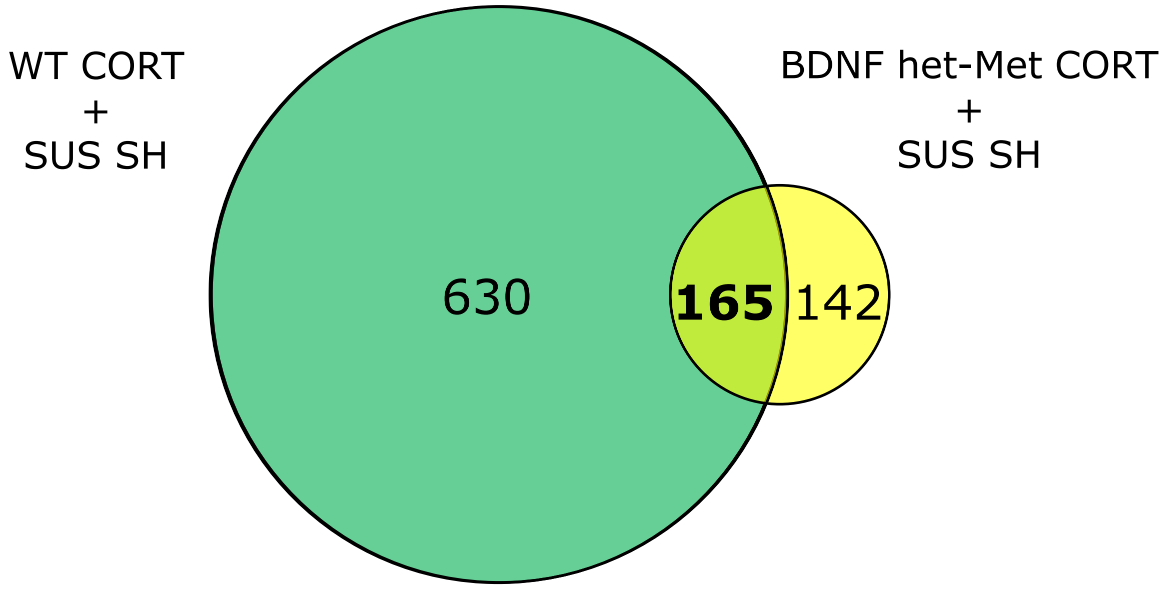


**Supplementary Fig. 1**

**Supplementary Table 1. Common GO and selected genes in RRHO selected quadrants**

| GO Term | Enrichment Score | # Genes | Select Genes |
|---|---|---|---|
| **Pink Module** | | | |
| Immunity | 16.21 | 73 | Fos, Ncf1, Ncf4, Iigp1, Tlr2, Tlr6, Casp8, Fcgr1, Nfkbia, H2-Aa |
| Extracellular Matrix | 10.90 | 53 | Cdh1, Cadm4, Itga1, Itgb2, Itgb3, Itgb4, Tgfbi, Des, Nectin2, Col2a1 |
| Wnt/PI3K Signaling | 9.70 | 90 | Wnt3, Wnt9a, Met, Hgf, Fgfr2, Tdgf1, Fgf18, Jak3, Tek, Comp |
| Signal Transduction | 5.11 | 159 | Grb7, Gab1, Igfbp2, Ghsr, Fyb, Hhip, Npbwr1, Pkig, Ntsr1, Oprk1 |
| Iron Homeostasis | 2.65 | 11 | Trf, Lcn2, Mfi2, Bdh2, Scara5, Steap1, Steap3, Steap4 |
| **Green Module** | | | |
| Neurotransmission | 11.98 | 64 | Cacna1b, Trpm2, Slc9a5, Cacna1g, Slc38a4, Clic6, Slc9a2, Scn5a, Kcnmb2, Casr |
| Neuronal Differentiation | 3.88 | 18 | Shh, Otx2, Notch1, Sox5, Ar, Kdr, Insm1, Kdm6b |
| Social Behavior | 2.53 | 6 | Tbx1, Nrxn2, Nrxn3, Shank1, Shank2, Cntnap2 |
| Vascular System | 2.52 | 7 | Cav3, Myh7, Hcn4, Casq2 |
| Phosphorylation | 1.46 | 68 | Myo1a, Myo5b, Smarca4, Tubd1, Mast1, Mast3, Mast4, Met, Dnm3, Fuk |

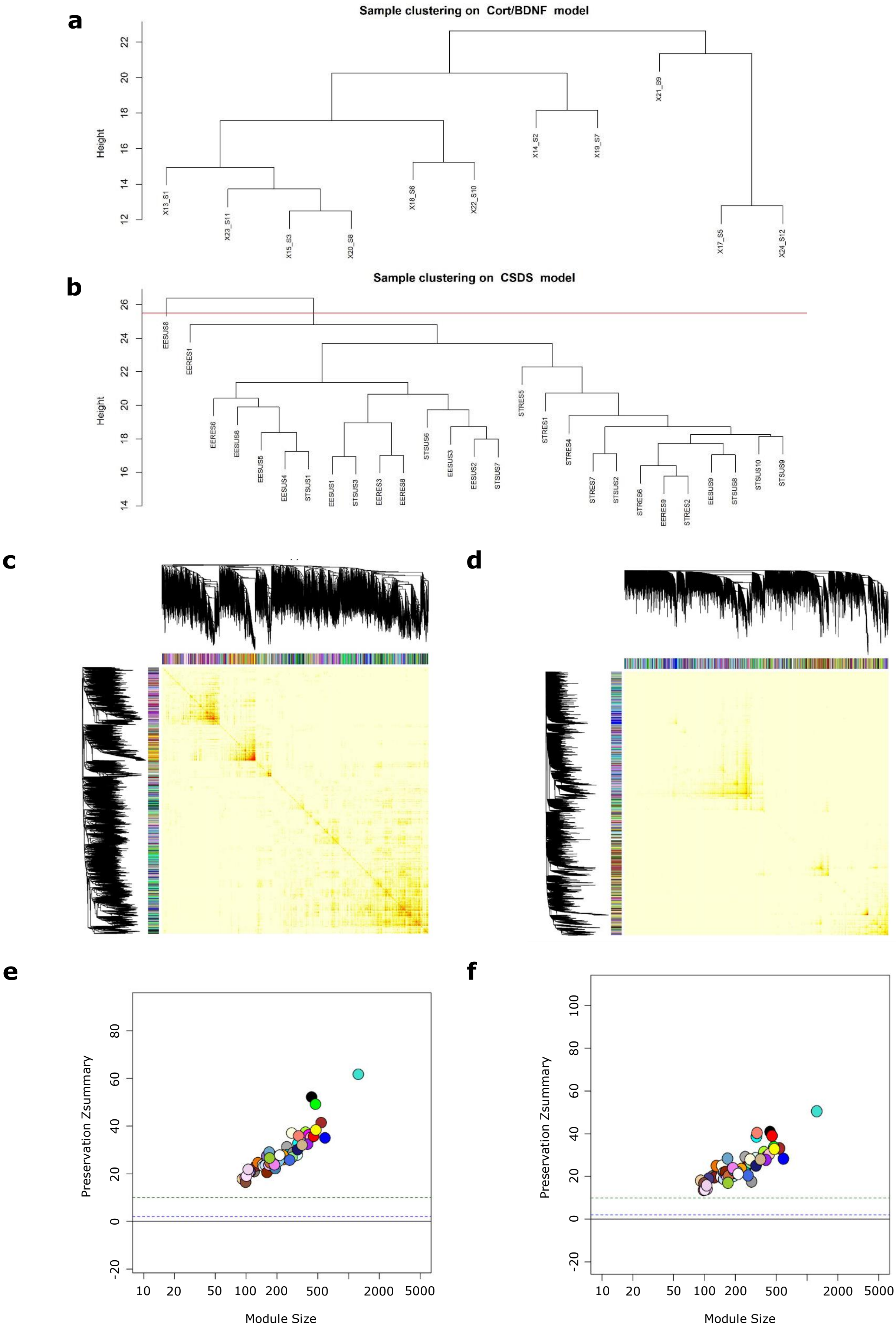


**Supplementary Fig. 2**

### SUPPLEMENTARY INFORMATION

**Supplementary Figure 1. Highly Ranked Genes across models** Venn diagrams showing common genes between experimental groups of the RRHO quadrants depicted in Fig. 2 corresponding to the point of highest overlap of each quadrant as described in Plaiser et al., 2010 (see Supplementary Methods). a) RES mice in SH and either WT mice (blue) or BDNF het-Met mice under vehicle (yellow), and between SUS in EE and either WT mice (purple) or BDNF het-Met mice under vehicle (coral), with 87 genes (Supplementary Data 2) shared across all comparisons (salmon); b) SUS mice in SH and either WT (green) or BDNF het-Met mice (yellow) under CORT, with 165 (Supplementary Data 2) shared across both comparisons (lime).

**Supplementary Table 1. Common GO and selected genes in RRHO selected quadrants** GO analysis was performed in RRHO quadrants identified to share genetic expression in similar biological networks. Quadrants with similar biological networks were aggregated to a "pink" module and a "green" module, as indicated in Figure 2. Table columns represent (i) Gene ontology term, (ii) enrichment score and (iii) total number of genes obtained from DAVID annotation clustering; (iv) genes selected based on their known role in neuronal function. Several of the most enriched GO terms were selected for representation.

**Supplementary Figure 2. Validation of Consensus Coexpression Network** Hierarchal clustering analysis for a) the model of CORT-treatment and b) the model of different housing conditions prior to chronic social defeat stress (CSDS). Modules with high mutual similarity of greater than 85% were merged. Topological overlap matrix (TOM) of WGCNA modules for c) the model of CORT-treatment and d) the model of different housing conditions prior to CSDS. Increasing color intensity from white to dark red within the TOM indicates increasing coexpression-based topological overlap. Preservation Z-summary of network modules as compared to their size in genes for e) the model of CORT treatment and f) the model of different housing conditions prior to CSDS. Modules with preservation Z-scores greater than 10 (above the dotted green line) and not less than 2 (below the dotted blue line) were indicated to be strongly preserved within the data set.
