## Supplementary material for "Genomic Modules and Intramodular Network Convergency of Susceptibility and Resilience in Multimodeled Stress in Male Mice": Supplementary Material Methods_FINAL.pdf

### Housing Conditions

To control for litter-specific effects[1, 2], mice in each group were selected from across multiple litters. CORT-treated animals were group housed (n=4–5) in standard cages (28.5x17x13cm) and were kept on a 12-h light-dark cycle (lights off 7:00pm) in a temperature-controlled room maintained at 21±2°C. A 2lux red light permitted animal maintenance in the dark phase. Food and water were available *ad libitum.* Enriched animals were housed in groups of 12 per 78x86cm cage, containing tunnels, novel inanimate objects, and running wheels. The enriched cages were fully cage changed every two weeks and a partial bedding change occurred once per week. Toys were cleaned once a week by soaking them for 10 minutes in PeroxyGard, then rinsing them in water, and then left to air dry. The position of the toys was changed weekly according to eight different configurations. Standard housed mice were grouped 3 per 30x18cm standard cage. The standard cages were cage changed every week. Based on previous behavioral studies, we found that a sample size of 8 to 12 mice per group allows us to reliably detect changes of the magnitude we are examining ($\alpha=0.05$). Variance is similar among the groups that are being compared. All procedures were performed in accordance with the National Guidelines on the Care and Use of Animals and a protocol approved by The Rockefeller University Animal Care and Use Committee.

### Chronic Oral CORT Treatment

At 2 months of age drinking water for WT or BDNF Val66Met mice was replaced with either a solution containing 25 µg/ml corticosterone (Sigma, St. Louis, MO) dissolved in 100% ethanol, and then diluted in regular tap water to a final concentration of 1% or a solution of 1% ethanol in tap water (vehicle). Both corticosterone and vehicle solutions were replaced twice a week.

### Chronic Social Defeat Stress (CSDS)

The mouse cohort raised in enriched versus standard housing underwent a modified social defeat stress procedure similar to that previously described[3] with some minor adjustments. Aggressor mice were first screened from retired CD-1 breeders. The CD-1 breeders at 4 months of age underwent an acclimatization period for 7 days in single housing with ad libitum access to food and water. Aggression screening occurs when a screener C57BL/6J mouse 8 to 11 weeks of age is placed in the CD-1 breeder's cage for 180 seconds then removed. The next day another novel screener is tested with the CD-1 mouse. This is then repeated for a final day. Criteria for selection of CD-1 mice for the social defeat protocol is recorded. This includes latency to the first attack and the total number of attacks for each CD-1 mouse. Mice that attack on at least two consecutive days in less than 60 seconds are included as aggressors. Mice that caused wounding in C57 mice during screening were excluded. The aggressors were ranked based on their aggression score (total sum of all attacks/total latency to the first attack). The CD1 aggressor with the highest aggression score after screening was also excluded. The social defeat stress occurred in a standard rat cage containing a Plexiglass divider with holes allowing for olfactory cues and separating the cage into two 1/3 and 2/3 sized compartments. A CD-1 aggressor was placed in the 2/3 size compartment and left overnight following its third screening session with access to food and water. The next day an intruder C57BL/6J mouse previously raised in enriched or standard housing was added to the 2/3 size compartment with the CD-1 aggressor. The two mice were left to interact for 5 minutes. If an attack bout occurs longer than 10 seconds the mice are separated momentarily. If the C57BL/6J mouse is wounded, the defeat session is terminated for the day. The amount of attacks a mouse received during the defeat session was also limited. After the defeat session, the C57BL/6J mouse is transferred to that cage’s 1/3 size compartment with adequate food and water. After 24 hours the C57BL/6J mouse is alternated to another cage of the same setup with a novel CD-1 aggressor where the social defeat procedure is repeated. This novel CD-1 aggressor is alternated between an aggressor with a high aggression score and an aggressor with a low aggression score. This pattern continues through the 10-day defeat protocol. This is an effort to maintain similar levels of total aggression exposure over the 10-day paradigm for each C57BL/6J mouse.

**Light-Dark Box Test**

The arena consisted of an open white-wall light box and a covered black-wall dark box (l=29cm, w=29cm). Mice were videotaped for 5 minutes by a camera fixed on the ceiling above the arena (50±10 lux). Time spent in the light box, latency to enter the dark box, and latency to reenter the light box were scored were scored by an experimenter blind to the experimental groups and conditions. Mice that did not enter the dark box during the 5-minute test cut-off were excluded from the analysis and from further investigation.

**Splash Test**

The splash test was performed as described by Santarelli et al.[4] and Isingrini et al.[5] with minor changes. After 30 minutes habituation to a new and empty testing cage, mice were then sprayed on the hindquarters with a 10% (w/v in water) sucrose solution, and immediately placed in the testing cage for 5 minutes (50±10 lux). Grooming behavior was videotaped by a camera fixed on the ceiling above the arena. The latency to the first grooming session, the total time grooming, and the number of grooming sessions were manually scored by an experimenter blind to the experimental groups and conditions. Animals that failed to perform grooming behavior during the 5-minute test cut-off were excluded from the analysis.

**Social Interaction Test**

After habituation, mice were placed into an open arena with an empty wire-mesh box at one side (interaction zone). Mice were given 150 s to explore the arena. A novel CD1 aggressor was placed in the box and the procedure was repeated. Time in the interaction zone, in the corner zones, and total locomotion were recorded by an experimenter blind to the experimental groups and conditions. Ratio of time inside and outside the interaction zone determined the classification of “resilient” (n=46) and “susceptible” (n=108). The ratio was calculated as =(time spent outside the interaction zone/time spent within the interaction zone).

**Behavioral Z-Score**

The z-score was calculated using the following formula: z=[(x-x̄cont)/σcont]*(±1)]. Z-scores across disparate models of stress were compared using a behavioral score for each animal is calculated as: 1- $z/\bar{a}$, where $z$=[(x-$\bar{x}_{cont}$)/$\sigma_{cont}$]*(±1)] and x=percentage time in the light, latency to dark, latency to light, grooming time, number of grooming sessions, grooming latency, time in social interaction zone, or SI ratio, and $a$=$z$ for each respective control mouse. After calculation, $a$ and $z$ values where normalized to the lowest $z$ score.

**RNA Sequencing**

Mice were sacrificed by cervical dislocation and rapidly decapitated to extract whole brain. The brains for CORT treated mice were immediately dissected to separate the ventral hippocampus, which was rapidly flash frozen and stored at -80 °C. CSDS mice whole brains were rapidly removed, flash frozen, and stored at -80 °C. CSDS brains were sliced coronally (200 μm) until bregma −2.30, then sliced horizontally 3.24 to 0.92 mm58. A 300 μm diameter puncher was used to punch the ventral dentate gyrus region. The RNA from CORT treated mice was extracted from the ventral hippocampal tissue using Qiagen Lipid Tissue Mini Kit (Qiagen, Germantown, MD USA). RNA from CSDS ventral dentate tissue was performed using Qiagen Allprep RNA Mini Kit (Qiagen, Germantown, MD USA). DNAse I treatment occurred during RNA extraction and RNA was examined by a Bioanalyzer (Agilent technologies, Santa Clara, USA) for both models. CORT sample cDNA libraries were sequenced on an Illumina NextSeq 500 to obtain single-end 75-bp reads at an approximate sequencing depth of 35-40 million reads per sample. RNA libraries from CSDS samples were prepared using Illumina TruSeq Stranded total

RNA LT set (Cat# RS-122-2301, Illumina Canada Ulc.) and sequenced on an Illumina HiSeq 2500, obtaining paired-end 125-bp reads at a depth of approximately 30 million reads per sample.

#### RRHO

The full threshold-free lists of differential expression data were first ranked by increasing log fold change. The RRHO2 “stratified” method (https://rdrr.io/github/RRHO2/RRHO2/) was used to detect the overlap between genes differentially expressed in the same or opposite directions[6], where the bottom right and top left quadrant display overlaps of genes with concordant differential expression, and the top right and bottom left discordant overlap. Threshold-free differential expression lists were ranked using the -log10(p-value) corresponding to the sign of the full change value generated from the limm-voom package. The point from each quadrant that has the highest absolute log10-transformed significance denotes the rank thresholds (and accordingly the metric thresholds) on the x- and y-axis (i.e. in both models) that would yield the most statistically significant set of overlapping differentially expressed genes or putatively co-regulated genes[7].

#### Gene Profile Processing

To fit gene co-expression network building by WGCNA, we first convert the raw reads count to RPKM (Reads Per Kilobase of transcript, per Million mapped reads) by using the formula: RPKM = numReads / (geneLength/1000 * totalNumReads/1,000,000). Meanwhile non-coding genes, genes in scaffold, and the genes that have more than 10% missing values in samples are filtered out. Clustering analysis also filtered out outliers. The hierarchical clustering function hclust was used to analyze the samples on their Euclidean distance with average linkage agglomeration method, separately in each set, which aims to filter out the outliers (See Supplementary Fig. 2a,b). The clustering height is the value of the criterion associated with the clustering method for the agglomeration.

#### Consensus Weighted Gene Co-Expression Network Construction

Gene coexpression analysis is useful in identifying transcriptional alterations in genetically complex disorders, where the phenotype is a consequence of numerous small genomic alterations rather than from isolated single-gene effects[8]. The consensus gene co-expression networks were constructed by the automatic block-wise network construction and module detection process in WGCNA package. The soft thresholding power $\beta$ (=18) to which co-expression similarity is raised to calculate adjacency was properly chosen based on the criterion of approximate scale-free topology before the consensus network building. The process firstly built the Topological Overlap Matrix (TOM) by calculating the gene adjacencies using biweight midcorrelation[9] with chosen soft thresholding power in the individual sets, then constructed the signed consensus weighted correlation network through scaling of individual TOM to make them comparable across sets, calculating consensus Topological Overlap, clustering and identifying module and merging of modules whose expression profiles are very similar. Modules were detected via hierarchical gene clustering on TOM-based dissimilarity and branch cutting using the top-down dynamic tree cut method[10]. Each Module could be represented by the module eigengene. Modules with high module eigengene correlations (r > 0.85) were merged. 54 modules, excluding grey, were identified.

**Module Preservation**

The module preservation analysis intends to evaluate whether the co-expression gene network is defined robustly in the current sample set and reproducible in independent data as well as whether the module is more significant than a random sample of genes[11]. $Z_{summary}$ statistics is calculated for module preservation analysis by combing module density-based statistics and intra-modular connectivity-based statistics and separability of modules based on permutation test ( Z=$\frac{observed-mean_{permutated}}{sd_{permuted}}$, $Z_{summary} = \frac{Z_{density}+Z_{connetivity}}{2}$). It has indicated that $Z_{summary}$< 2 indicates no preservation, 2 < $Z_{summary}$ < 10 weak to moderate evidence of preservation, and $Z_{summary}$ > 10 strong evidence of preservation. We performed permutation 100 time to reconstruct the networks with the same parameters with randomly resampling the initial sample set. The module preservation analysis results for each individual model are shown in Supplementary Fig. 2e,f.

**KME Measurement for Hub Genes**

In generating a weighted gene co-expression network, each module can be represented by the module eigengene. The module eigengene turns out to be the most highly connected gene. Eigengene-based connectivity, also known as kME is simply the correlation between the genes expression profile and the module eigengene[9].

$$K_{ME,i} = Cor(X_i, ME)$$

We calculated the consensus KME value and the corresponding P-value.

### Gene Network of Hub Genes in Significant Modules

To investigate the connectivity of the hub genes, ARACNE (Algorithm for the Reconstruction of Accurate Cellular Networks)[12] was used first to identify the significant interactions between the hub genes in the significant modules based on their mutual information. For each ARACNE-derived unweighted network, the connectivity scores of the hub genes were computed. The hub genes which are associated with enriched biological processes were examined to identify if the hub genes were enriched in brain function related GO.

### REFERENCES


1. Becker G, Kowall M. Crucial role of the postnatal maternal environment in the expression of prenatal stress effects in the male rats. J Comp Physiol Psychol. 1977;91(6):1432-46.
2. Chapman RH, Stern JM. Failure of severe maternal stress or ACTH during pregnancy to affect emotionality of male rat offspring: implications of litter effects for prenatal studies. Dev Psychobiol. 1979;12(3):255-67.
3. Golden SA, Covington HE, 3rd, Berton O, Russo SJ. A standardized protocol for repeated social defeat stress in mice. Nat Protoc. 2011;6(8):1183-91.
4. Santarelli L, Saxe M, Gross C, Surget A, Battaglia F, Dulawa S, et al. Requirement of hippocampal neurogenesis for the behavioral effects of antidepressants. Science. 2003;301(5634):805-9.
5. Isingrini E, Camus V, Le Guisquet AM, Pingaud M, Devers S, Belzung C. Association between repeated unpredictable chronic mild stress (UCMS) procedures with a high fat diet: a model of fluoxetine resistance in mice. PLoS One. 2010;5(4):e10404.
6. Cahill KM, Huo Z, Tseng GC, Logan RW, Seney ML. Improved identification of concordant and discordant gene expression signatures using an updated rank-rank hypergeometric overlap approach. Sci Rep. 2018;8(1):9588.
7. Plaisier SB, Taschereau R, Wong JA, Graeber TG. Rank-rank hypergeometric overlap: identification of statistically significant overlap between gene-expression signatures. Nucleic Acids Res. 2010;38(17):e169.
8. Gaiteri C, Ding Y, French B, Tseng GC, Sibille E. Beyond modules and hubs: the potential of gene coexpression networks for investigating molecular mechanisms of complex brain disorders. Genes Brain Behav. 2014;13(1):13-24.
9. Langfelder P, Horvath S. Fast R Functions for Robust Correlations and Hierarchical Clustering. J Stat Softw. 2012;46(11).
10. Langfelder P, Horvath S. WGCNA: an R package for weighted correlation network analysis. BMC Bioinformatics. 2008;9:559.
11. Langfelder P, Luo R, Oldham MC, Horvath S. Is my network module preserved and reproducible? PLoS Comput Biol. 2011;7(1):e1001057.
12. Meyer PE, Lafitte F, Bontempi G. minet: A R/Bioconductor package for inferring large transcriptional networks using mutual information. BMC Bioinformatics. 2008;9:461.
